# The proSAAS chaperone provides neuroprotection and attenuates transsynaptic α–synuclein spread in rodent models of Parkinson’s disease

**DOI:** 10.1101/2021.09.29.462435

**Authors:** Iris Lindberg, Zhan Shu, Hoa Lam, Michael Helwig, Nur Yucer, Alexander Laperle, Clive Svendsen, Donato A. Di Monte, Nigel T. Maidment

## Abstract

Parkinson’s disease is a devastating motor disorder involving the aberrant aggregation of the synaptic protein synuclein (aSyn) and degeneration of the nigrostriatal dopaminergic tract. We previously showed that proSAAS, a small secreted chaperone protein widely expressed in neurons within the brain, is able to block aSyn-induced dopaminergic cytotoxicity in primary nigral neuron cultures. We show here that coinjection of proSAAS-encoding lentivirus profoundly reduced the motor asymmetry caused by unilateral nigral AAV-mediated human aSyn overexpression. This positive functional outcome was accompanied by significant amelioration of the human aSyn-induced loss of both nigral tyrosine hydroxylase-positive cells and striatal tyrosine hydroxylase-positive terminals, demonstrating clear proSAAS-mediated protection of the nigro-striatal tract. ProSAAS overexpression also reduced the content of human aSyn protein in both the nigra and striatum and reduced the loss of tyrosine hydroxylase protein in both regions. Since proSAAS is a secreted protein, we tested the possibility that proSAAS is able to block the transsynaptic spread of aSyn from the periphery to the central nervous system, increasingly recognized as a potentially significant pathological mechanism. The number of human aSyn-positive neurites in the pons and caudal midbrain of mice following administration of human aSyn-encoding AAV into the vagus nerve was considerably reduced in mice coinjected with proSAAS-encoding AAV, supporting proSAAS-mediated blockade of transsynaptic aSyn transmission. We suggest that proSAAS may represent a promising target for therapeutic development in Parkinson’s disease.

**Significance:** This paper describes two independent avenues of research that both provide support for the *in vivo* neuroprotective function of this small chaperone protein. In the first approach, we show that proSAAS overexpression provides remarkably effective protection against dopaminergic neurotoxicity in a rat model of Parkinson’s disease. This conclusion is supported both by three independent assays of motor function as well as by quantitative analysis of surviving dopaminergic neurons in brain areas involved in the control of motor function. In the second line of research, we show that in mice, the spread of human synuclein across synapses can be blunted by proSAAS overexpression.

## INTRODUCTION

Neurodegenerative diseases such as Alzheimer’s disease (AD) and Parkinson’s disease (PD) are increasingly considered to arise, at least in part, from the cumulative effects of abnormal aggregating proteins. Environmental toxins and other stressors accumulated during aging disrupt normal proteostatic mechanisms, resulting in the accumulation of misfolded and aggregated proteins. Cellular chaperones represent an important component of the neuronal proteostasis network that works both to combat initial protein misfolding and to direct misfolded proteins towards degradative pathways. Consistent with this idea, overexpression of chaperone proteins has been found to be cytoprotective against toxic protein aggregates in animal models of a variety of neurodegenerative diseases (reviewed in (1-3)). Specific examples include cytoprotection by overexpression of heat shock proteins in fly and rodent models of PD (4-7); reviewed in (8). In addition, disaggregase-type chaperones, such as the yeast disaggregase Hsp104 (9) and the mammalian disaggregase Hsp110 (10) have been shown to block alpha-synuclein (aSyn) pathology and cytotoxicity. The beneficial effects of chaperone overexpression on brain proteostasis are associated with improvements in both motor and cognitive function (reviewed in (2, 11, 12)).

While *cytoplasmic* chaperones such as heat shock proteins have been frequently studied in the context of neurodegenerative disease, relatively little attention has been paid to the possible contribution of *secreted* brain chaperone proteins to brain proteostasis. Secreted chaperones, such as the ubiquitously-expressed glycoprotein clusterin (also known as ApoJ), are uniquely positioned to intercept and/or to blunt the toxicity of extracellular aggregates both prior to and following endocytic cell entry (reviewed in (13)). However, as yet there are few *in vivo* studies showing rescue of neurodegenerative defects by secreted chaperones other than clusterin (reviewed in (14)). Thus the question of whether other secreted chaperones play a role in blocking the cytotoxicity and/or the pathogenic transmission of aggregating proteins has not been adequately addressed.

Given the devastating consequence of proteostatic dysregulation in the CNS, and the apparent ability of pathogenic aggregating proteins to travel throughout the brain (reviewed in (15)) it would not be surprising to find that neurons express specific secreted chaperones to combat the formation and propagation of pathogenic aggregates. Of the few known secreted chaperones preferentially expressed in brain, the small protein known as proSAAS (encoded by the gene *PCSK1N*) exhibits many of the features expected of such a chaperone. ProSAAS is expressed in neurons rather than glia (16), where it is stored within secretory vesicles (17, 18) and presumably released into the synaptic cleft, well positioned to function as an extracellular chaperone at the synapse. Our past work has shown that recombinant proSAAS blocks the aggregation of Abeta 1-42 and aSyn at highly sub-stoichiometric ratios (19). More importantly, both proSAAS overexpression as well as exogenous application of recombinant proSAAS are able to reduce Abeta 1-42 and aSyn-mediated neurotoxicity, both in neuronal cell lines and in primary cultured neurons (19, 20). However, whether proSAAS overexpression is able to functionally protect neurons *in vivo* has yet to be established. Here we demonstrate the profound behavioral and neuropathological protective effect of proSAAS overexpression in a virus-mediated nigral human aSyn overexpression rat model of Parkinson’s disease. Additionally we show that proSAAS overexpression decreases the pathogenic transmission of human aSyn in a mouse model of aSyn brain spreading following injection of viral vectors expressing human aSyn into the vagus nerve.

## RESULTS

### ProSAAS overexpression protects rats from human aSyn-induced motor asymmetry

In order to test the protective potential of proSAAS overexpression on human aSyn-induced motor dysfunction, rats received unilateral nigral injections of either proSAAS-encoding or GFP-encoding lentivirus together with AAV-encoding human aSyn, a treatment known to produce degeneration of nigrostriatal neurons. Additional control groups of rats received either GFP-encoding lentivirus plus GFP-encoding AAV, or PBS. Rats were subjected to a weekly battery of tests 2 weeks prior to, and 8 weeks following injections to assess motor asymmetry. Of note, this fully controlled experiment was preceded by an independent pilot experiment - using a different batch of proSAAS-expressing lentivirus -that produced similar evidence of both the behavioral- and neuroprotective effect of proSAAS overexpression.

**Figure 1** shows data from each of the 3 motor asymmetry tests conducted – cylinder test (**Figure 1A**), bracing test (**Figure 1B**), and forelimb placement test (**Figure 1C**). Profound motor asymmetry in the human aSyn + GFP group is apparent in all 3 tests, the progressive nature of which is reflected in a significant group by time interaction using a 2-way repeated measures ANOVA: cylinder test F(27, 304) = 3.800, p < 0.0001; bracing test F(27, 306) = 38.26, p < 0.0001; forelimb placement test F(27, 304) = 27.90, p < 0.0001. In each case, this motor asymmetry was largely prevented by co-injection of proSAAS-encoding virus. For the cylinder test, the preference for use of the forepaw ipsilateral to the injection evident in the aSyn + GFP group was significantly curtailed in the aSyn + proSAAS-treated animals (main effect of group F_(3,34)_ = 13.52, *p* < 0.0001; post-hoc Tukey comparison aSyn + GFP vs. aSyn + proSAAS, *p* < 0.0001). Similarly, for the bracing test, the reduced number of adjustment steps contralateral to the injected side relative to the ipsilateral forepaw, apparent in the aSyn + GFP group, was largely absent in the aSyn + proSAAS-treated animals (main effect of group F_(3,34)_ = 74.23, *p* < 0.0001; post-hoc Tukey comparison aSyn + GFP vs. aSyn + proSAAS, *p* < 0.0001). Finally, in the forelimb placement test, the reduction in number of forelimb placements contralateral to the injection relative to ipsilateral in aSyn + GFP-treated animals was significantly curtailed by co-expression of proSAAS (main effect of group F_(3,34)_ = 40.98, *p* < 0.0001; Tukey comparison aSyn + GFP vs. aSyn + proSAAS, *p* < 0.0001). Taken together, these data provide clear evidence that overexpression of proSAAS provides substantial protection from aSyn-induced motor asymmetry.

**FIG. 1.**
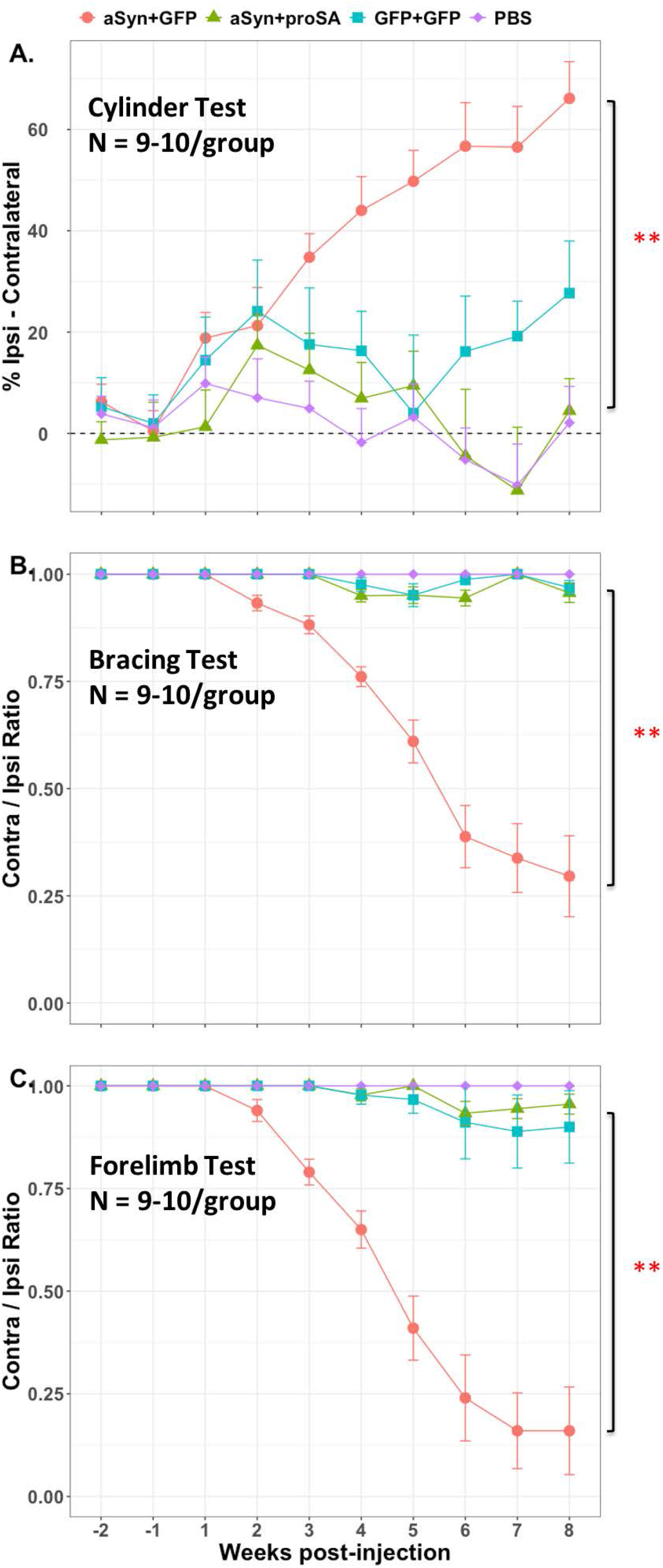
ProSAAS overexpression prevents development of motor asymmetry induced by nigral human aSyn expression. Animals (9-10 per group) received unilateral nigral injections of combinations of AAV and lentivirus expressing human aSyn, proSAAS and GFP, or PBS, as described in the text. Motor asymmetry was assessed by 3 tests, each conducted at weekly intervals beginning 2 weeks before the injection. Each point represents the mean ± SEM of asymmetry scores. Progressive motor asymmetry evident in rats injected with human aSyn-expressing AAV + GFP-expressing lentivirus was not observed in rats injected with human aSyn-expressing AAV + proSAAS-expressing lentivirus. (A) **Cylinder test** (Injection: F_(3,34)_ = 13.52, *p* < 0.0001). (B) **Bracing test** (Injection: F_(3,34)_ = 74.23, *p* < 0.0001). (C**) Forelimb placement test** (Injection: F_(3,34)_ = 40.98, *p* < 0.0001) \*\**p* < 0.0001 post-hoc Tukey, aSyn+GFP vs. aSyn+proSAAS.

### ProSAAS overexpression attenuates human aSyn-induced nigrostriatal DA neuron degeneration

To determine if the functional protection provided by proSAAS overexpression was accompanied by sparing of the nigrostriatal tract from aSyn-induced damage we examined brain slices through the SNc and striatum immunostained for tyrosine hydroxylase (TH). Representative sections from each of the 4 groups for which behavioral data are presented above are shown in **Figure 2A**, demonstrating clear asymmetry in TH immunoreactivity in both the SNc and striatum in the example from the aSyn + GFP group, which is markedly less apparent in the examples from the 3 other groups.

**FIG. 2.**
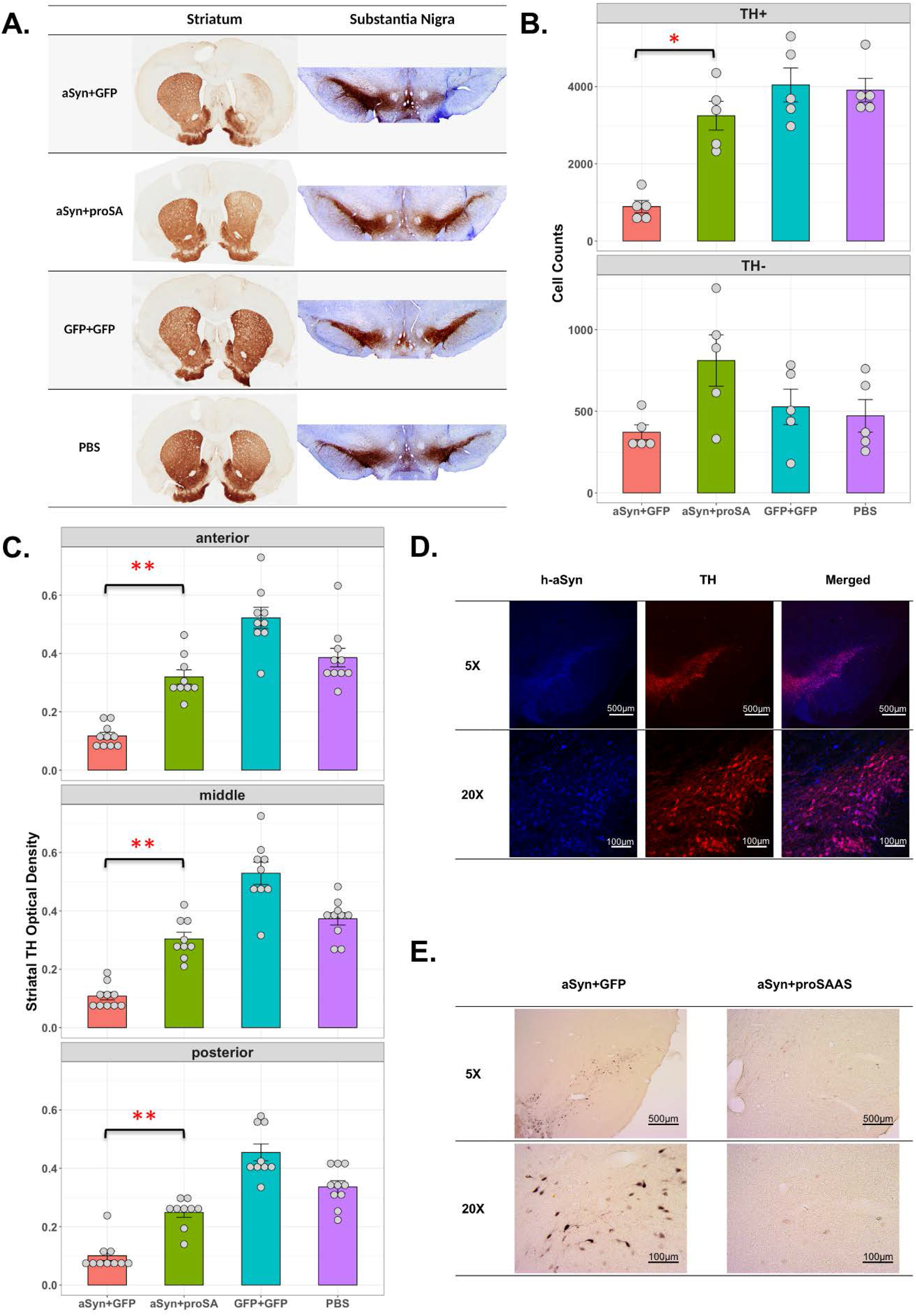
ProSAAS overexpression protects nigro-striatal dopaminergic neurons from human aSyn-induced toxicity. (A) Representative immunohistochemically-stained brain slices containing striatum and substantia nigra. (B) Number of TH-positive (TH+) and TH negative (TH-) neurons in the SNc ipsilateral to the injection site measured by unbiased stereology (N = 5/group). ProSAAS preserved TH+ neurons in the SNc (**TH+**: one-way ANOVA, main effect of group: F_(3,16)_ = 19.03, *p* < 0.0001; post-hoc Tukey, aSyn+GFP vs. aSyn+proSAAS: \**p* < 0.001. **TH-**: one-way ANOVA, F_(3,16)_ = 2.93, *p* > 0.05). (C) TH optical density measured at 3 striatal rostro-caudal levels ipsilateral to the injection site: anterior, middle and posterior (N = 9-10/group). Two-way ANOVA, main effect of group: F_(3,34)_ = 57.81, *p* < 0.0001; main effect of striatal level: F_(2,68)_ = 14.42, *p* < 0.0001; Interaction: F_(6,68)_ = 1.243, *p* = 0.296). ProSAAS protected TH immunoreactivity in the striatum (post-hoc Tukey, aSyn+GFP vs. aSyn+proSAAS: \*\**p* < 0.0001 in all segments). (D) Representative section through the SNc from an aSyn+proSAAS-treated rat showing localization of human aSyn in TH-positive cells. (E) Similarly representative SNc sections showing the pronounced labeling with anti-p129 aSyn Ab in aSyn+GFP, but not aSyn+proSAAS-treated rats.

Quantitation of this apparent neuroprotective effect of proSAAS overexpression was conducted by unbiased stereological counting of nigral TH-positive cells in the SNc and by TH densitometry in the striatum. These measures were made bilaterally; however, to avoid potential confounding effects of compensatory changes contralateral to the injection site, statistical analyses were restricted to between-group comparisons of the sites ipsilateral to the injection (**Figure 2**). Full bilateral data are presented in **Supplementary Figures S1 and S2. Figure 2B** (upper panel) shows the profound reduction in number of TH-positive cells in the SNc of rats injected with human aSyn - plus GFP-encoding viruses relative to GFP + GFP and PBS groups, and the considerable attenuation of this reduction in the human aSyn + proSAAS group (main group effect F_(3,16)_ = 19.03, *p* < 0.0001; post-hoc Tukey comparison aSyn + GFP vs. aSyn + proSAAS, *p* < 0.001). Importantly, the number of Nissl-positive, TH-negative cells did not differ significantly between the 4 groups (lower panel) indicating that the reduction in TH-positive cells in the aSyn + GFP group was due to degeneration of DA cells rather than down-regulation of TH expression in otherwise intact neurons.

Quantitation of striatal terminal TH density was conducted at 3 rostro-caudal levels, designated anterior, middle and posterior, and is shown in **Figure 2C**. Statistical analysis of optical density by 2-way ANOVA revealed a main effect of group (F_(3,34)_ = 57.81, *p* < 0.0001), a main effect of rostro-caudal segment (F_(2,68)_= 14.42, *p* < 0.0001), but no group × segment interaction (F_(6,68)_ = 1.243, *p* = 0.296). Similar to nigral cell counts, striatal terminal density was significantly lower in human aSyn + GFP-treated animals compared to animals in the human aSyn + proSAAS group (post-hoc Tukey, *p* < 0.0001). Moreover, the protection offered by proSAAS was such that TH terminal density did not differ significantly from that of PBS controls (post-hoc Tukey, *p* > 0.05). Notably, however, there remained a significant difference from the GFP + GFP group, which itself was significantly higher than the other three groups (post-hoc Tukey, *p* < 0.0001 in each comparison). Further, this apparent increase in TH density was bilateral (**Supplementary Figure 2**). We conclude that lentiviral proSAAS overexpression provides significant protection from human aSyn-induced neuronal damage to the nigro-striatal DA tract.

### aSyn phosphorylation in the SNc is reduced in proSAAS-treated animals

Colocalization of human aSyn and TH in surviving neurons in the SNc of animals treated with proSAAS was readily apparent (**Figure 2D**). Importantly, however, aggregation-prone phosphorylated aSyn staining (31-35) was considerably reduced by proSAAS treatment. **Figure 2E** shows representative examples of extensive p129 aSyn staining in the SNc of animals treated with AAV-aSyn + lenti-GFP 8 weeks post-injection, and minimal staining in animals treated with AAV-aSyn + lenti-proSAAS. No such staining was apparent in sections from GFP/GFP-treated animals.

### ProSAAS overexpression reduces virus-produced human aSyn protein levels in SNc and striatum and attenuates the reduction in TH protein 3 weeks post-treatment

Since one function of chaperones is to promote the degradation of toxic aggregating proteins, we sought evidence for reduced protein levels of human aSyn in the SNc and striatum of animals injected with proSAAS- and human aSyn-encoding viruses relative to those receiving aSyn-plus GFP-expressing viruses. This was assessed by Western blotting 3 weeks after unilateral injection, a time of predicted maximal viral expression but before significant cell loss was expected. A GFP-GFP control group was included to demonstrate specificity, and examples of gels from each of the 3 groups are shown in **Figure 3A**. Quantitation of blot intensities, normalized to β-actin loading controls, are shown in **Figure 3B**. Statistical analysis of intensities on the injected (right) side across striatum (combined dorsal and ventral caudate signals) and SNc indicated a significantly lower level of human aSyn protein in animals injected with human aSyn plus proSAAS virus relative to those injected with aSyn plus GFP virus (ANOVA of aligned rank transformed data, F_(1,7)_ = 9.24, *p* = 0.019). Further, analysis of TH protein levels revealed significant attenuation of human aSyn-induced reduction in TH protein by co-expression of proSAAS (main effect of treatment F_(2,10)_ = 22.09, *p* = 0.0002; post-hoc p<0.005). Bilateral quantitation data are provided in **Supplementary Figure S3**. Human aSyn expression was restricted to the injected side, as expected. Interestingly, however, TH protein levels were lower in aSyn + GFP treated animals relative to GFP + GFP animals on both sides of the brain, perhaps reflecting a compensatory mechanism, or a significant contribution of down-regulation in crossed nigro-striatal neurons. This effect of aSyn contralateral to the site of injection was also attenuated by unilateral proSAAS co-administration such that considerable asymmetry in TH levels persisted.

**FIG. 3.**
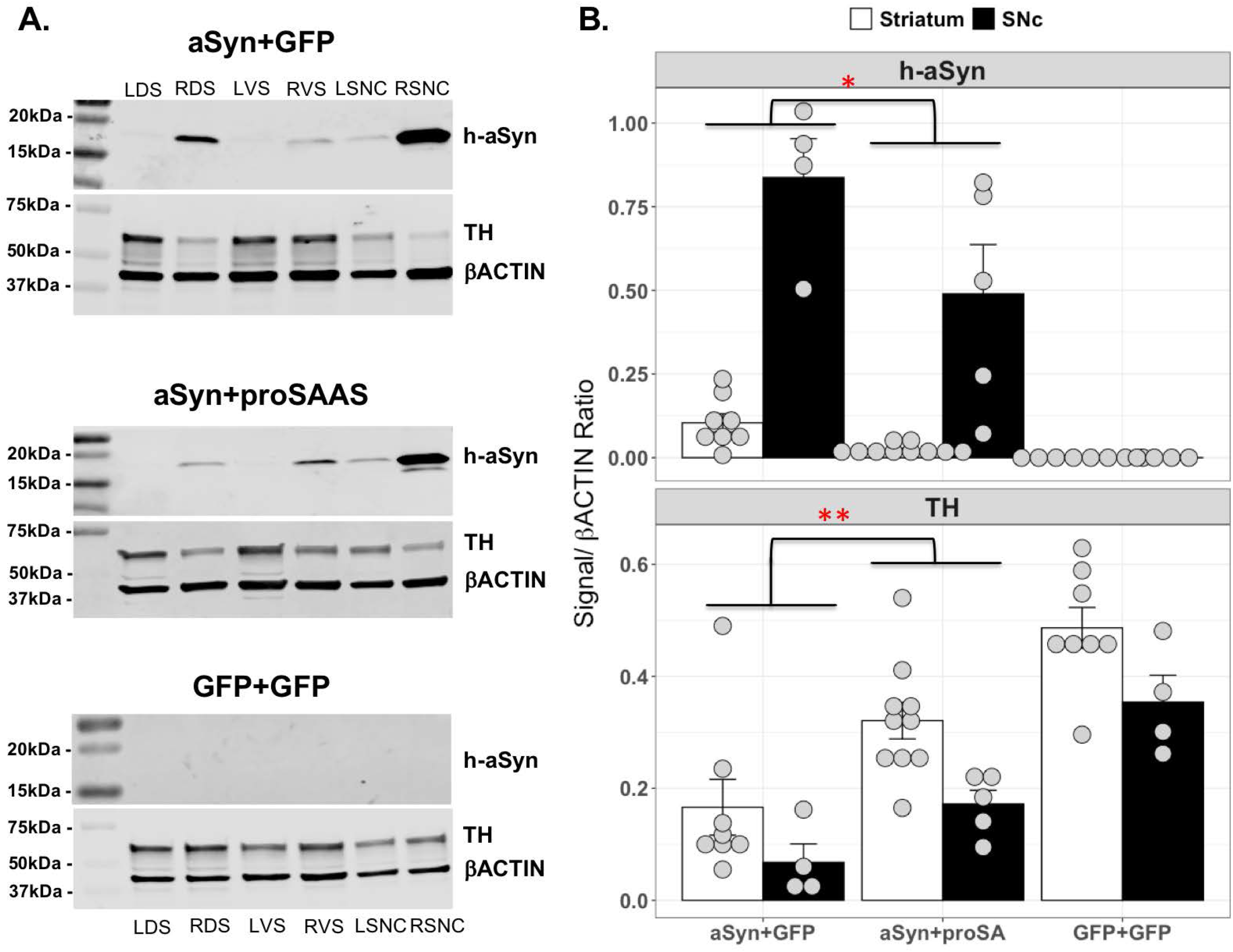
ProSAAS overexpression reduces human aSyn protein levels and attenuates the drop in TH protein levels in nigrostriatal tissues. (A) Representative membranes containing human aSyn (18 kDa), TH (60 kDa) and beta-actin (42 kDa) blots (LDS: left dorsal striatum; RDS: right dorsal striatum; LVS: left ventral striatum; RVS: right ventral striatum; LSNC: left substantia nigra pars compacta; RSNC: right substantia nigra pars compacta; left: contralateral side; right: ipsilateral side). (B) Expression levels of human aSyn and TH normalized to their beta actin levels on the injected (right) side of the striatum (dorsal and ventral combined), and the SNc (N = 4-5 /group). The level of human aSyn protein was significantly reduced across these regions (ANOVA of aligned rank transformed data, aSyn + proSAAS vs. aSyn + GFP: F(1,7) = 9.24, *p = 0.019). Further, human aSyn-induced reduction in TH protein levels was attenuated by simultaneous proSAAS expression. (Two-way ANOVA with repeated measurements, treatment: F(2,10) = 22.09, p = 0.0002; post-hoc Tukey, aSyn + proSAAS vs. aSyn + GFP: **, p < 0.005).

### ProSAAS expression attenuates the caudo-rostral transmission of human aSyn in mice

In order to determine whether proSAAS expression can block the transmission of aSyn, we injected human aSyn-encoding AAV together with either proSAAS- or GFP-encoding AAV unilaterally into the vagus nerve of mice (**Figure 4**). This experimental strategy was designed to specifically overexpress human aSyn in the medulla oblongata (MO) and then to follow its temporal spreading into rostral regions of the brain. As expected, asymmetric human aSyn immunoreactivity (more intense ipsilateral to the vagal injection site) was observed 6 weeks later in neurons and dendrites of the MO, as shown by representative examples in the top two panels of **Figure 4**, where **panel A** represents human aSyn-immunoreactivity in an animal injected with aSyn AAV plus proSAAS AAV, while **panel B** represents a similar image from a mouse injected with aSyn AAV plus GFP-encoding AAV.

**FIG. 4.**
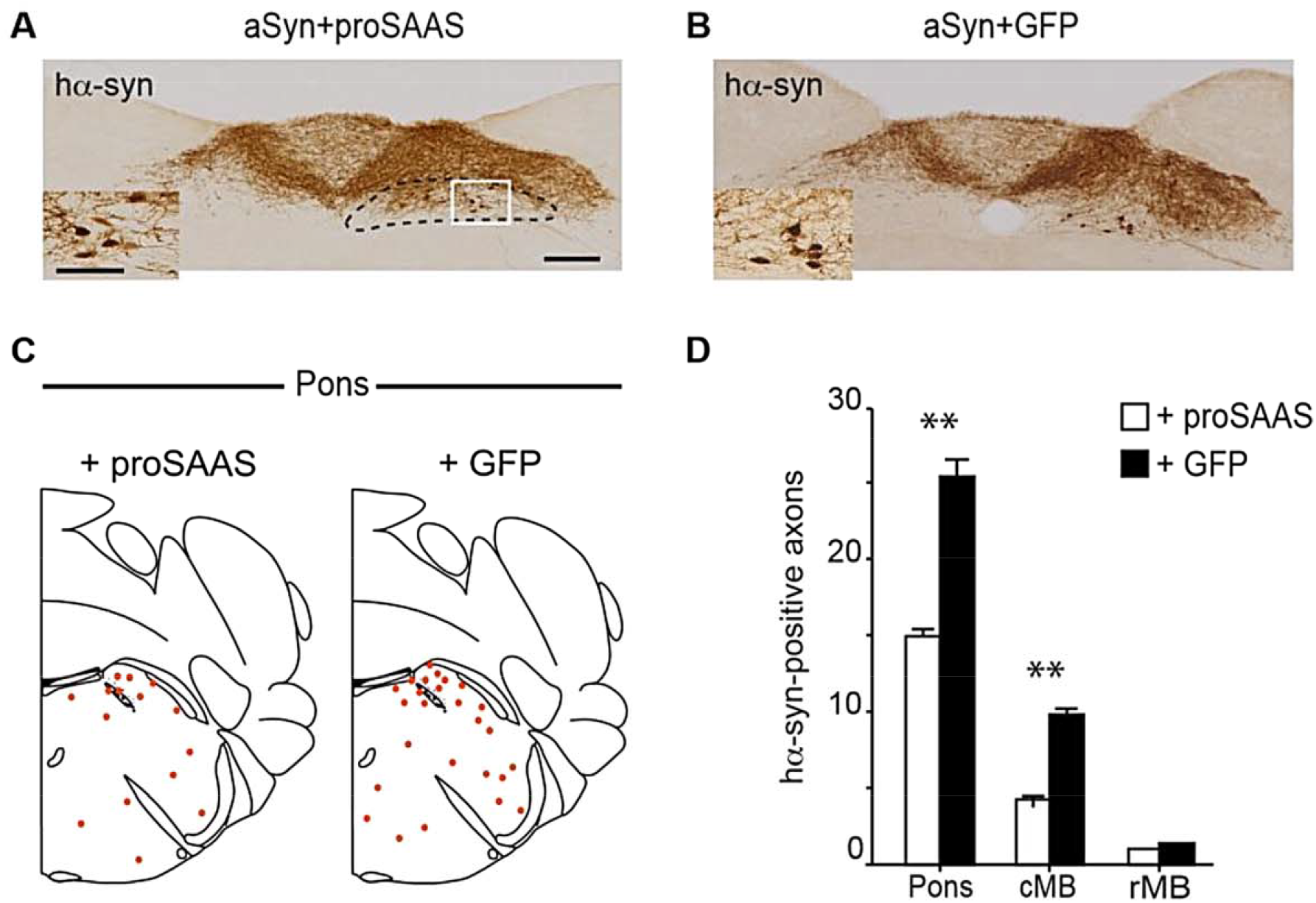
ProSAAS overexpression attenuates caudo-rostral brain spreading of aSyn in mice. Mice were sacrificed 6 weeks after a single injection into the left vagus nerve of a cocktail containing either human aSyn + proSAAS-expressing AAVs **(A)**; or human aSyn + GFP-expressing AAVs **(B). Panels A** and **B** are representative sections through the dorsal medulla oblongata from these 2 groups, immunostained with anti-human aSyn. The dorsal motor nucleus of the vagus (DMnX) is delineated by a dashed line, and the square boxes indicate the area shown at higher magnification. *Scale bars*: Low magnification = 200 μ m; high magnification = 50 μ m. (**C**) Schematic representation of the distribution of human aSyn-immunoreactive axons in proSAAS/aSyn-versus GFP/aSyn-co-expressing mice. (**D**) Counts of human aSyn-immunoreactive axons in the pons, caudal midbrain (cMB), and rostral midbrain (rMB) of AAV-injected mice (n = 5/group). **, p= 0.0025 (pons) or 0.0083 (cMB).

Similar to the observations in the rat SNc and striatum (**Figure 3**), the intensity of aSyn immunoreactivity in the mouse MO was reduced in the proSAAS-co-expressing group (scores: +, n = 1; ++, n = 3; +++, n = 1) compared to the GFP-co-expressing group (scores: +++, n = 5). This reduction in the MO was accompanied by decreased numbers of human aSyn-positive-axons in the pons and caudal midbrain. Thus, trans-synaptic spread from the medulla oblongata was significantly reduced in proSAAS-treated mice relative to their GFP-treated counterparts (**Figure 4C and D)**. No such diminishment was apparent in the rostral midbrain (rMB), but it should be noted that aSyn transmission had only minimally spread this far in controls. These data replicate a prior experiment and support the idea that proSAAS overexpression attenuates the trans-synaptic spread of aSyn through the brain.

## DISCUSSION

The data presented here showing that proSAAS exerts strong protection from aSyn-induced nigro-striatal DA tract degeneration provide the first *in vivo* evidence of the potential utility of this chaperone as a therapeutic target for PD. AAV-mediated overexpression of human wildtype aSyn unilaterally in the rat SNc produced a robust progressive motor asymmetry over an 8-week period accompanied by significant loss of TH-positive nigral cells and TH-positive striatal terminals measured post-mortem, consistent with previous reports describing similar viral-mediated aSyn models (reviewed in (36)). All three of these measures of aSyn toxicity were profoundly blunted by nigral co-injection of proSAAS-expressing lentivirus. Further, the rostral spread of aSyn in the brain of mice following vagal injection of AAV-expressing human aSyn was similarly attenuated by co-vagal injection of AAV-proSAAS suggesting that therapeutic strategies targeting proSAAS may have potential for slowing progression of PD.

ProSAAS, a small neuronal and endocrine protein first identified in an unbiased peptidomics screen from brain extracts over 20 years ago (37), has now been identified by ten independent proteomics groups as a consistently lowered CSF biomarker in AD as well as in frontotemporal dementia (see meta-analysis in (38); see also (39, 40)); one recent report also indicates lower proSAAS levels in PD CSF (39). Immunoreactive proSAAS is associated with Lewy bodies in the brains of PD patients (19), with amyloid plaques in AD patients (20), and with tau tangles in various forms of dementia (41); reviewed in (14), suggesting brain sequestration of proSAAS during disease progression. In agreement with this idea, transcriptomics studies of AD patient tissues indicate increased brain proSAAS RNA expression during AD progression (42). Increased brain proSAAS is also seen in patients with cerebral amyloid angiopathy (43) as well as in proteomics studies of two animal models of neurodegenerative disease, equine serum sickness (44) and a rat model of Usher’s syndrome (45). In rodents, hypothalamic levels of proSAAS increase following stressors including dehydration, hypoxia, and cold temperature (46-48), and recent work from our laboratory has shown that proSAAS levels are upregulated in Neuro2A cells during tunicamycin-induced endoplasmic reticulum stress (49). Collectively, these data, obtained from a variety of human disease and cell and animal models, support the idea that proSAAS is a stress-responsive protein that contributes to brain proteostasis during the progression of neurodegenerative disease.

aSyn, an abundant brain protein, is thought to participate in synaptic vesicle recycling and may have additional functions at the synapse (reviewed in (50)). Aggregation of aSyn into Lewy bodies, a key signature of Parkinson’s disease, is believed to result in the loss of normal aSyn function as well as a gain of toxic function, leading to general synaptic dysfunction and ultimately neuronal cytotoxicity (51); reviewed in (52). The relative susceptibility of DA neurons to the toxic effect of aSyn overexpression and aggregation remains a subject of debate but may involve a role for DA or its metabolites in the aggregative process itself and/or a general elevated level of oxidative stress imparted by DA metabolism, which may be heightened when synaptic cycling and therefore vesicle packaging of DA is disrupted (53); see (54, 55) for review. The precise cellular mechanism underlying the strong cytoprotective action of proSAAS observed in the current *in vivo* study and in our previous cellular investigation (19) is not yet clear, but most likely depends upon its demonstrated chaperone activity (reviewed in (14)). If proSAAS were acting simply as a secreted trophic factor to promote neuronal health (56), we might expect transsynaptic transmission of aSyn to be increased in the presence of increased proSAAS; instead, we found diminished transsynaptic spread of this protein.

As noted in the Introduction, proSAAS is one of only a few identified brain-specific *secretory* chaperones. The protective effect of *cytoplasmic* heat shock protein overexpression in animal models of PD has been confirmed in multiple studies (reviewed in (14)). Effective reversal of cytotoxic, and in some cases, motor deficits in PD rodent models by cytoplasmic chaperone overexpression has been accomplished using Hsp70 (5, 6); Hsc70 (7); the Hsp70-interacting protein BAG1 (57); and the disaggregases Hsp104 and Hsp110 (9, 10). Most recently, the ubiquitin ligase TRIM11 has been shown to possess aSyn chaperone activity and can rescue aSyn-mediated cell death in cell and animal models of PD (58). These chaperones are thought to exert their cytoprotective actions by directly blocking the intracellular formation of toxic aSyn aggregates, providing strong support for the idea that chaperone overexpression can restore normal proteostasis.

While it is clear that proSAAS is a highly potent anti-aggregant attenuating aSyn fibrillation (19), its sequestration to the secretory pathway limits its presence in the cytoplasm to endosomal uptake. While the cytoplasm clearly represents the predominant cellular location of aSyn aggregates, a small portion of cytoplasmic aSyn is known to be secreted (reviewed in (15)), indicating a possible extracellular location of interaction with proSAAS. Thus proSAAS may exert its effects on aSyn both within the synapse and following endosomal reuptake. These actions could take several forms. For example, secreted proSAAS, upon binding to extracellular aSyn, could block the uptake of aSyn into neurons (and/or other cells such as astrocytes). Indeed, our observation that proSAAS overexpression attenuates the spread of aSyn through the brain would be consistent with such a mechanism. ProSAAS may also exert cytoplasmic chaperone action following reuptake of proSAAS-bound toxic oligomers, promoting the direction of toxic aSyn aggregates to degradative pathways. In this case, functional interaction of proSAAS with aSyn could take place both in the extracellular space and in the cytoplasm. While the Western blot data presented above appear to support a facilitating effect of proSAAS on aSyn degradation, it is also possible that they instead reflect the blockade by proSAAS of the assembly of toxic oligomeric species, which again, could be initiated either extracellularly or intracellularly. Finally, given the association of aSyn with synaptic vesicle cycling and the important protective role that vesicular packaging of DA is believed to play in dopaminergic neurons, proSAAS’ neuroprotective efficacy may lie not only in its general ability to prevent the buildup of toxic aSyn assemblies, but also in its ability to maintain normal aSyn function at its primary site of action in the synaptic vesicle cycling machinery.

In conclusion, the data presented here indicate that local manipulation of brain proSAAS levels exerts a strong protective effect on nigrostriatal DA neurons against direct aSyn-mediated toxicity; and further, that increasing proSAAS levels within the vagus nerve limits the spread throughout the brain of similarly targeted aSyn overexpression. We suggest that these findings offer promise for the utility of PD therapies targeting proSAAS in halting the progression of PD, perhaps even following peripheral routes of intervention.

## MATERIALS AND METHODS

### Neuroprotection study in rats

#### Subjects

Male Sprague-Dawley rats (Charles River, USA) weighing 250-320 grams at the time of surgery were maintained on a 12-12 h light-dark cycle. Food and water were available *ad libitum* throughout the study. Animal care conformed to the U.S. Public Health Service Guide for the Care and Use of Laboratory Animals, and procedures were approved by UCLA Institutional Animal Care and Use Committee.

#### Viruses

Adeno-associated viral vectors (AAV 2/1) expressing human aSyn or GFP, driven by the chicken beta actin promoter and enhanced using a woodchuck hepatitis virus post-transcriptional regulatory element (WPRE), were obtained from the UNC Viral Core at titers of 9.8 and 9.7 × 10^12^ viral genome copies (gc)/ml, respectively. Lentiviruses expressing mouse *Pcsk1n* (proSAAS) or eGFP, driven by the CMV promoter, were synthesized by GeneCopoiea (Rockville, MD) in the Lv153 backbone at titers of 4.7 × 10^8^ and 4.5 x10^8^ TU/ml, respectively.

#### Virus Injection

Surgeries were carried out under isoflurane anesthesia (5% in oxygen for induction and 2% for maintenance; Zoetis, Kalamazoo, MI). Pre-emptive analgesia was provided by an injection of Carprofen (5 mg/kg s.c.) prior to surgery. During surgery, animals were secured in a stereotaxic frame, with fascia retracted, skull revealed, and four burr holes drilled at the following coordinates above the substantia nigra pars compacta (SNc): AP −4.7 mm, ML +2.1 mm; AP −5.2 mm, ML +1.7 mm; AP −5.6 mm, ML +1.3 mm; AP −5.6 mm, ML +2.4 mm. A second dose of Carprofen was administered 24 h after the initial dose.

Animals were randomly assigned to 4 groups, each receiving 4 unilateral injections into the right SNc of a mixture of two viruses or phosphate buffered saline (PBS) vehicle as follows: 1) AAV2/1-human aSyn + lentivirus-proSAAS (n=9), 2) AAV2/1-human aSyn + lentivirus-GFP (n=10), 3) AAV2/1-GFP + lentivirus-GFP (n=9), 4) PBS (n=10). Each injection was 2 µl in volume and was delivered into the SNc at a rate of 0.2 µl/min, using the above-mentioned coordinates at a depth of −8.4 mm, −8.6mm, −8.6mm and −7.6mm, respectively, via a 30-gauge 10 µL Hamilton syringe (Hamilton 701SN-30/3’/3) mounted on a motorized injector (Stoelting QSI). After each injection, the syringe remained *in situ* for a further 5 min before withdrawal.

A separate cohort of rats was injected with combinations of viruses as described above (4-5 per group) and euthanized 3 weeks later for Western analysis of nigral and striatal aSyn and tyrosine hydroxylase (TH) protein content as described below.

#### Motor asymmetry tests

A battery of three tests known to be sensitive to dopamine (DA) depletion, without the need for extensive training, drug administration, or food or water deprivation, was employed, as previously described (21). All animals were handled and habituated to the behavioral testing procedures for 4 days and baseline scores on each test were subsequently obtained once weekly for two weeks before virus injection. Weekly testing resumed one week after surgery and continued for eight weeks. Tests were scored manually without knowledge of group assignment. The *cylinder test* examines spontaneous forelimb utilization during vertical movements. Rats were placed in a clear Plexiglas cylinder (diameter = 20 cm, height = 30 cm) and video-recorded over a 5-min period. During rearing movements, placement of a single paw onto the cylinder wall was recorded as an ipsilateral or contralateral placement relative to the injected side. When both paws were placed on the cylinder simultaneously or in rapid succession, both paws remaining on the cylinder, a score of ‘both’ was recorded. An asymmetry score was calculated as the difference between ipsilateral (right) and contralateral (left) forepaw placements as a percentage of total placements. The *bracing test* probes the ability of the rat to adjust postural balance in response to examiner-imposed lateral movement. The rat was held with only one limb (a forelimb) unrestrained to support its weight and moved laterally at a constant speed across a flat surface for 1 meter in 5 seconds. The number of adjusted steps was recorded twice for both leftward movements with the left forepaw and rightward movements with the right forepaw. Test scores were calculated by dividing the number of steps contralateral to the injected side (leftward movements) by the number of ipsilateral (rightward) steps. The *forelimb placement* test examines sensorimotor/proprioceptive capacity. During the test, the rat’s torso was held with both forelimbs hanging freely and moved slowly sideways toward a vertical flat surface until the vibrissae of one side touched the surface, evoking a characteristic placement of the adjacent forepaw on the surface. Tests of both ipsilateral and contralateral sides were repeated 10 times. The score was calculated as the ratio between the number of successful placements of the contralateral and ipsilateral forelimbs.

#### Post-mortem histological assessment

##### Tissue preparation

Following the final motor asymmetry test, animals were anesthetized with pentobarbital (100 mg/kg) and transcardially perfused with PBS followed by 4% paraformaldehyde in PBS. Brains were harvested, cryoprotected in 30% sucrose at 4°C overnight, and stored at −80°C. Brains were cut coronally on a cryotome (Leica CM1850) at 40 µm, and sections washed three times in 1x phosphate-buffered saline (PBS) to remove cryoprotectant, incubated in 0.3% H_2_O_2_ for 15 min, and washed again in PBS. Sections were then blocked with 5% normal goat serum/0.5% Triton X-100 in PBS for 2 h at room temperature, followed by overnight incubation at 4°C with a primary antibody for tyrosine hydroxylase (TH) (rabbit, 1:1000, Millipore #657012) diluted in blocking solution. The next day, sections were washed 3 times in 3X PBS and once in PBS (10 min/wash). Sections were incubated with a secondary antibody (goat anti-rabbit 1:300, Vector Labs #BA-1000) diluted in 2% normal goat serum/0.1% Tween for 2 h. The sections were washed in 0.1% Tween/PBS and then incubated for 1 h in 0.1% Tween/PBS solution containing an avidin-biotin complex (Vectastain, Vector Labs). After subsequent washing in 0.1% Tween/PBS, the sections were developed using diaminobenzidine (DAB, Sigma). Brain sections were mounted on glass slides separated by regions of interest (striatum or SNc) and left to air dry. Nissl staining was performed 24 h after immunostaining by rehydrating the sections in water. Tissue sections were then incubated in a filtered Cresyl violet solution for 10 min, followed by a quick wash, and immersed in 95% ethanol/0.1% glacial acetic acid. Sections were dehydrated in increasing concentrations of ethanol, soaked in xylene, and mounted with Eukitt mounting medium.

##### Unbiased stereology

Cell counting was performed by an investigator blinded to the virus injection group using Stereo Investigator (MBF Bioscience) coupled with a Leica DM-LB microscope and a Ludl XYZ motorized stage. Five animals from each group were randomly selected from each group. The absolute number of neurons in the SNc was estimated using the optical fractionator method (22). One in every 8 sections was used to estimate the number of neurons in the entire SNc in the virus-injected hemisphere and in the contralateral hemisphere. The counting frame was 50 × 50 × 20 µm (length × width × dissector height), and the sampling grid was 150 × 150 µm with a 3 µm guard zone at the top and bottom of the section. The average final section thickness was calculated from the measurements at each counting frame. Following delineation of the SNc under a 5X objective, counting was performed with a 100X oil objective. Cell morphology revealed by Nissl staining was used for classification and identification of different cell types in the SNc using criteria detailed in (23-26). Cells with a visible nucleus stained for TH immunoreactivity were counted as TH-positive neurons. Cells stained with Cresyl violet with a heavily-stained large single nucleus and lacking TH staining were counted as TH-negative neurons.

##### Striatal TH optical density assessment

Immunostained, mounted, striatal sections were scanned using an Aperio ScanScope at 10 µm resolution and the striata ipsilateral and contralateral to the injection analyzed with ImageJ software (version 1.53). Boundaries of the striatum at three anterior-posterior levels relative to Bregma – anterior +2.16 mm, middle +1.08 mm, posterior −0.05 mm - were determined by comparing anatomical landmarks in sections with a rat brain atlas (27).

##### Immunocytochemical localization of virally expressed human aSyn in the SNc

Midbrain sections from perfused brains removed from rats injected unilaterally in the SNc with human aSyn-expressing AAV plus proSAAS-expressing lentivirus were processed as described above. Sections were first washed 3 times for 10 min each with PBS then incubated with 10mM citric acid and 0.05% Tween 20 for 10 min at 90°C. Following cooling, sections were washed 3 times for 10 min/each with PBS before incubating in 5% donkey serum (Jackson ImmunoLab), 1% BSA and 0.5% Triton X-100 in PBS for 2 h at room temperature with shaking. Sections were then incubated in primary antibodies overnight at 4°C: sheep anti-TH (Millipore, #AB1542) at 1:1000 dilution, mouse anti-human aSyn (Life Technologies, #180215) at 1:200 dilution. Sections were washed 3 times with 3X PBS for 10 min/each and 1 time with PBS and incubated for 2 h at room temperature with secondary antibodies - Alexa Fluor 555 donkey anti-sheep at 1:1000 dilution (Invitrogen, cat# A21436), Alexa Fluor 647 donkey anti-mouse at 1:200 dilution (Invitrogen, #A31571) in 2% donkey serum and 0.1% Tween in PBS. After washing 3 times with 0.1% Tween/PBS and once with PBS, sections were mounted and cover-slipped prior to visualization under a microscope and image capture. Some sections were alternatively processed for phosphorylated aSyn imaging. Such sections were washed as described above, followed by immersion in H_2_O_2_ for 15 min, and subsequently washed a further three times in PBS. Sections were then blocked with 5% normal goat serum/0.5% Triton X-100/1% BSA in PBS for 2 h at room temperature, followed by overnight incubation with a primary rabbit anti-aSyn phospho-129 antibody (1:5000, Abcam #51253) diluted in the same blocking solution. The next day, sections were washed 3 times in 3X PBS and once in PBS (10 min/wash). Sections were incubated with a secondary antibody (goat anti-rabbit 1:300, Vector Labs #BA-1000) diluted in 2% normal goat serum/0.1% Tween for 2 h. The sections were washed in 0.1% Tween/PBS and then incubated for 1 h in 0.1% Tween/PBS solution containing an avidin-biotin complex (Vectastain, Vector Labs). After subsequent washing in 0.1% Tween/PBS, the sections were developed using diaminobenzidine (DAB, Sigma). Brain sections were mounted on glass slides for microscopic analysis and image capture.

#### Western analysis of nigral and striatal aSyn and TH protein levels

Fresh-frozen brains were stored at −80 and subsequently mounted in a cryostat. Tissue from the SNc was collected from 4 × 300 μm sections using a 1.5 mm tissue punch. Similarly, tissue from the striatum (dorsal and ventral caudate, separately) was collected from 6 × 300 μm sections using 2 mm and 1.5 mm tissue punches, respectively. Tissues were disrupted using a 1 ml pipet tip and lysed using 1X NETN buffer (20 mM Tris-HCl, pH 8.0, 100 mM NaCl, 0.5 mM EDTA and 0.5% NP-40) supplemented with phosphatase/protease inhibitor cocktail (MS-SAFE, Sigma-Aldrich) in a 4°C water bath using a sonicator (QSONICA, Q800R3) for 20 min. Samples were centrifuged for 20 min at 4°C at 15000 rpm. Total soluble protein concentrations were measured using a Bradford assay (BIO-RAD). Concentrated Laemmli sample buffer (4x, BIO-RAD #161-0774) was added to extracts containing 50 µg of total protein and samples were boiled for 10 min. Samples were electrophoresed on 4-20% Mini-PROTEAN TGX Precast gels (BIO-RAD, 456-1094) and transferred to PVDF membranes using a Trans-Blot Turbo Transfer System (BIO-RAD). Membranes were blocked with Odyssey blocking buffer (LI-COR) and then incubated with primary antibodies overnight at 4°C. Following incubation with dye-labeled secondary antibodies for 2 h at room temperature, signals were imaged using an Odyssey Fc imaging system (LI-COR). Primary Western blotting antibodies were: human aSyn (1:1000, Abcam #ab138501), TH (1:2000, ImmunoStar, #22941), and β-actin (1:5000, Sigma-Aldrich, #A5441). Secondary Western blot antibodies were IRDye 680 RD goat anti-mouse LI-COR 926-68070), and IRDye 800CW goat anti-rabbit, (LI-COR 926-32211), all used at a dilution of 1:5000. The comparison of aSyn and TH protein levels was made following normalization of bands to β-actin.

#### Statistical analyses

Comparisons of motor asymmetry scores collected over multiple weeks were analyzed by ANOVA corrected for repeated measures. Group comparisons of post-mortem data were analyzed by ANOVA followed by a Tukey *post hoc* test when data were normally distributed. When the presumption of normal distribution was not justified, a Kruskal-Wallis test was performed followed by a Mann-Whitney U test with Bonferroni correction. Western blot data were analyzed using ANOVA with aligned rank transform followed by the least-squares means with Tukey corrections. All statistical analyses were performed with custom R code (R Project for Statistical Computing – http://www.R-project.org/).

### Transsynaptic aSyn spread study in mice

#### Subjects

Experiments were carried out on 12-week-old female C57BL/6JRj mice (Janvier). Experimental protocols were approved by the State Agency for Nature, Environment and Consumer Protection in North Rhine Westphalia. Mice were maintained on a 12-12 h light-dark cycle. Food and water were available *ad libitum* throughout the study.

#### Viruses

AAV-mediated transgene expression of human aSyn (AAV 2/6; Sirion Biotech), proSAAS (AAV 2/6; Vigene Biosciences) or enhanced green fluorescent protein (GFP; AAV 2/6; Vector Biolabs) was driven by the human SYN1 promoter, enhanced using a woodchuck hepatitis virus post-transcriptional regulatory element and a polyadenylation signal sequence (28, 29). Stock preparations were diluted to generate injection titers of 8 × 10^12^ gc/ml for human aSyn-AAV and 4 × 10^13^gc/ml for both proSAAS- and GFP-AAVs.

#### Surgical procedure

Following anesthetization with 2% isoflurane mixed with O_2_ and N_2_O, a 1-cm incision was made at the midline of the neck. The left vagus nerve was isolated from the carotid artery, and vector solution (800 nl) was injected at a flow rate of 350 nl/min using a 35-gauge blunt steel needle fitted onto a 10 ml NanoFil syringe as described previously (30). Mice received a mixture of either aSyn-AAV + proSAAS-AAV (n=5) or aSyn-AAV + GFP-AAV (n=5). After injection, the needle was kept in place for two additional minutes. Post-surgery analgesia was provided by subcutaneous injection with buprenorphine (Temgesic, 0.1 mg/kg). Six weeks after injection, mice were euthanized under pentobarbital anesthesia by perfusion through the ascending aorta first with saline containing heparin and then with ice-cold 4% (w/v) paraformaldehyde.

#### Post-mortem histological assessment

##### Tissue preparation

Brains were removed, immersion-fixed in 4% paraformaldehyde and cryopreserved in 30% (w/v) sucrose solution. Coronal sections (35 μm) throughout the brain were cut using a cryostat and stored at −20°C in phosphate buffer (pH 7.4) containing 30% glycerol and 30% ethylene glycol.

##### Brightfield microscopy and quantitative analyses

Free-floating brain sections were processed for brightfield microscopy as previously described (29). Primary antibody was directed against rabbit anti-h-aSyn (ab138501, MJFR1, Abcam; 1:50,000). Sections were rinsed and incubated for 1 h at room temperature in biotinylated secondary antibody solution (Vector Laboratories; BA1000, 1:200). Following treatment with avidin-biotin-horseradish peroxidase complex (PK 6100; ABC Elite kit, Vector Laboratories), the color reaction was developed using a 3,3′-diaminobenzidine kit (Vector Laboratories). Images were obtained using an Observer.Z1 microscope (Carl Zeiss) equipped with a motorized stage. Low magnification overview images were generated with a 20× Plan-Apochromat objective (N.A. 0.8) followed by computerized image stitching with ZEN 2 software (Carl Zeiss). For z-stack high-magnification imaging, stacks were collected at 2 μm intervals with a 63× Plan-Apochromat N.A. 1.4) objective using the same system, followed by deep focus postprocessing. Staining of aSyn in the DMV and NTS at the level of the medulla oblongata (MO, Bregma: Bregma −7.48 mm), was scored into categories of + (low), ++ (medium) and high (+++) intensities by two independent blinded observers.

The number of human aSyn-positive-axons was counted in sections at the level of the pons (Bregma: - 5.40 mm), caudal midbrain (cMB; Bregma: −4.60 mm) and rostral midbrain (rMB, Bregma: −3.40 mm) using an Axioscope microscope (Carl Zeiss) under a 63x Plan-Apochromat objective. Scoring and counting were performed by investigators blinded to the treatment protocol. Data were analyzed using an unpaired Student’s t test and are shown as the mean + SEM (n= 5 fields).

## Acknowledgements

We are grateful to R. Rusconi for construction of the proSAAS AAV2 shuttle vector. This work was supported by NIH grant AG 062222 to I. Lindberg and N.T. Maidment.

## Abbreviations

AAV: adeno-associated virus
aSyn: alpha synuclein
cMB: caudal midbrain
DA: dopamine
DMV: dorsal motor nucleus of the vagus
GFP: green fluorescent protein
MO: medulla oblongata
NTS: nucleus tractus solitarius
rMB: rostral midbrain
SNc: substantia nigra pars compacta
TH: tyrosine hydroxylase

## Supplementary Figure Legends

**Supplementary Figure S1:**
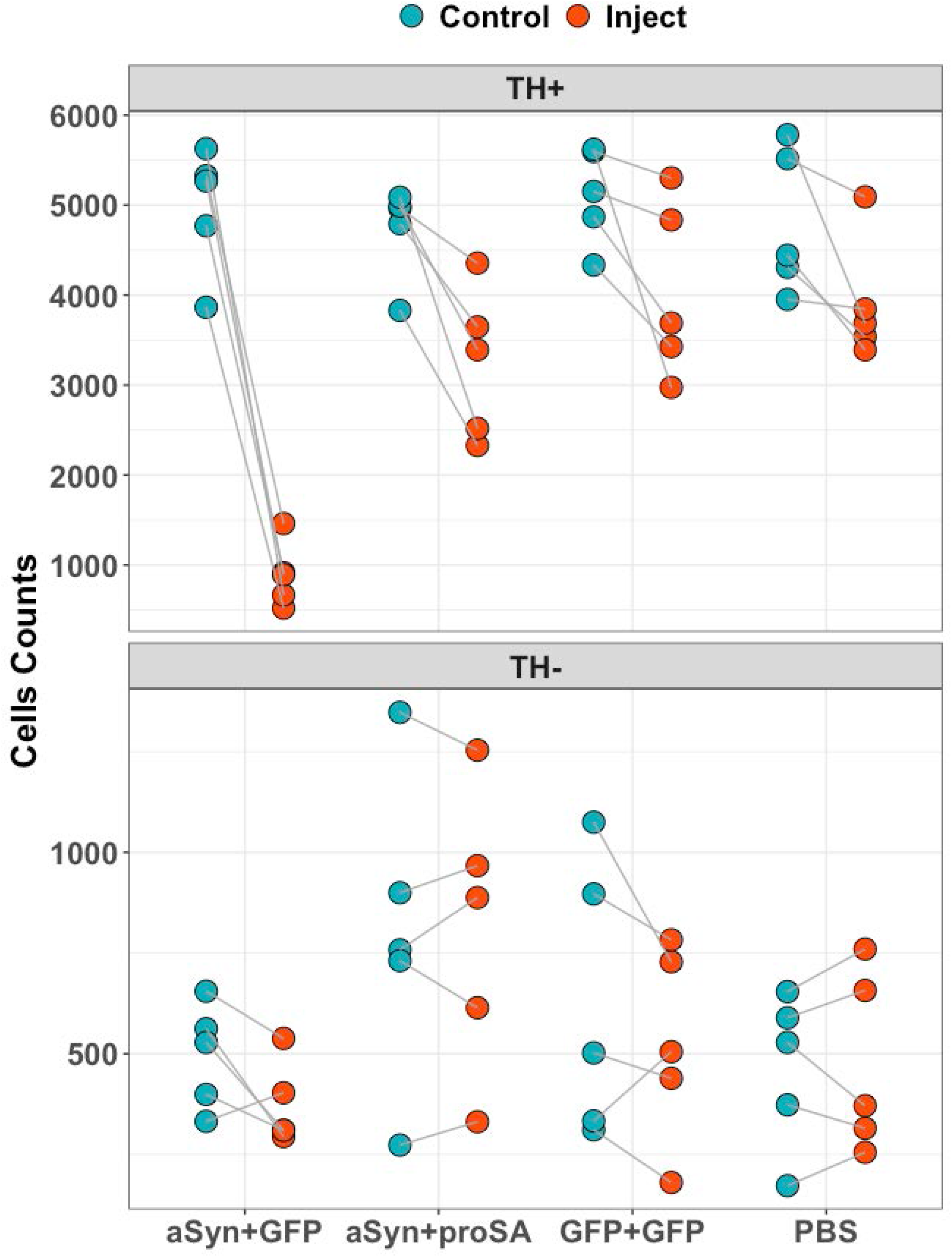
ProSAAS protects nigral dopaminergic neurons from human aSyn-induced toxicity. Bilateral stereological assessment of TH-positive and TH-negative neurons in the SNc demonstrating asymmetry in the number of TH-positive neurons, control versus injected side, in animals injected with aSyn + GFP viruses, which is attenuated in animals injected with a combination of aSyn- and proSAAS-expressing viruses.

**Supplementary Figure S2:**
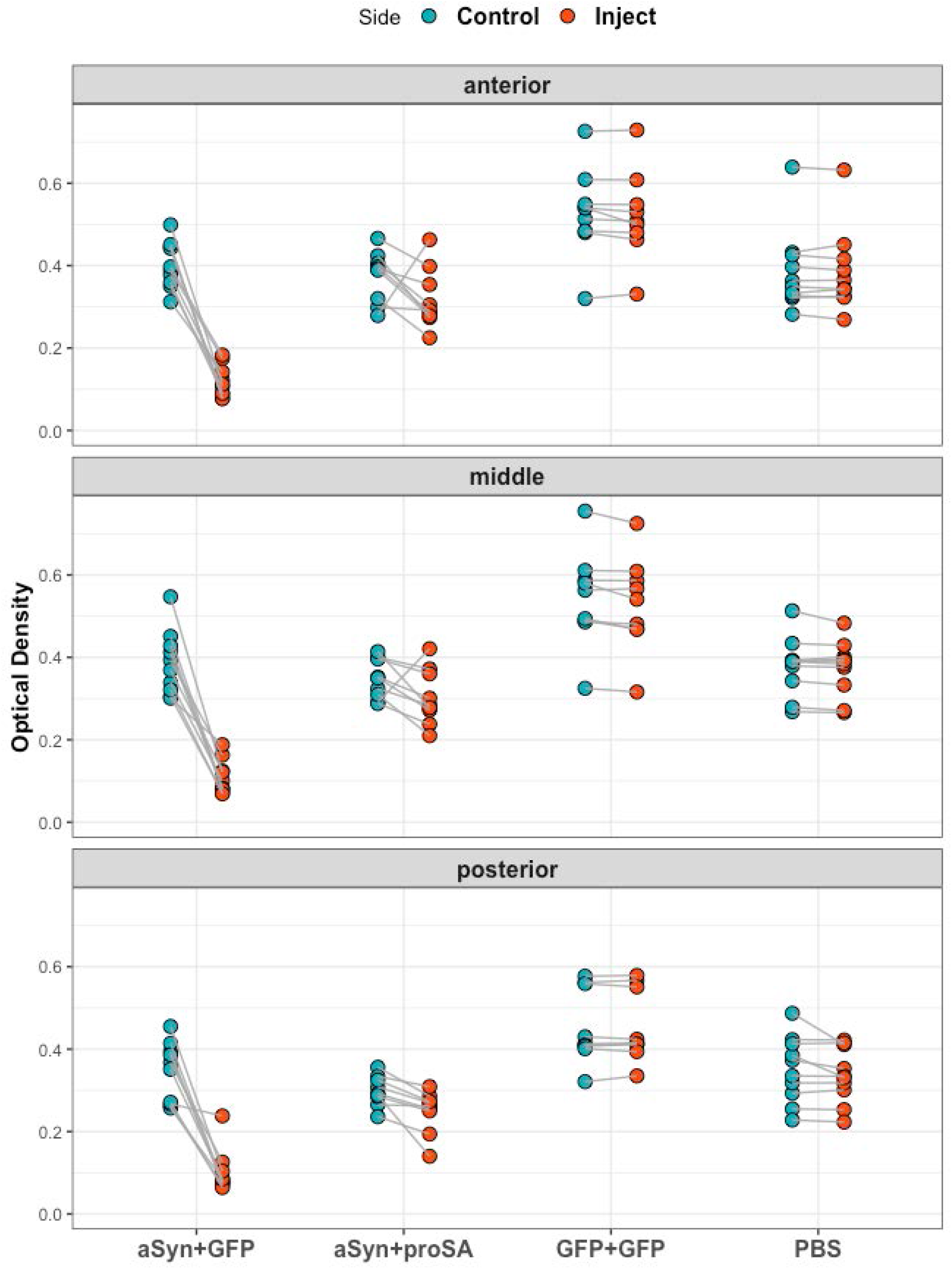
ProSAAS protects striatal dopamine terminals from human aSyn-induced toxicity. Bilateral assessment of striatal TH immunoreactivity by densitometry demonstrating asymmetry, control versus injected side, in animals injected with aSyn + GFP viruses, which is attenuated in animals injected with a combination of aSyn- and proSAAS-expressing viruses.

**Supplementary Figure S3:**
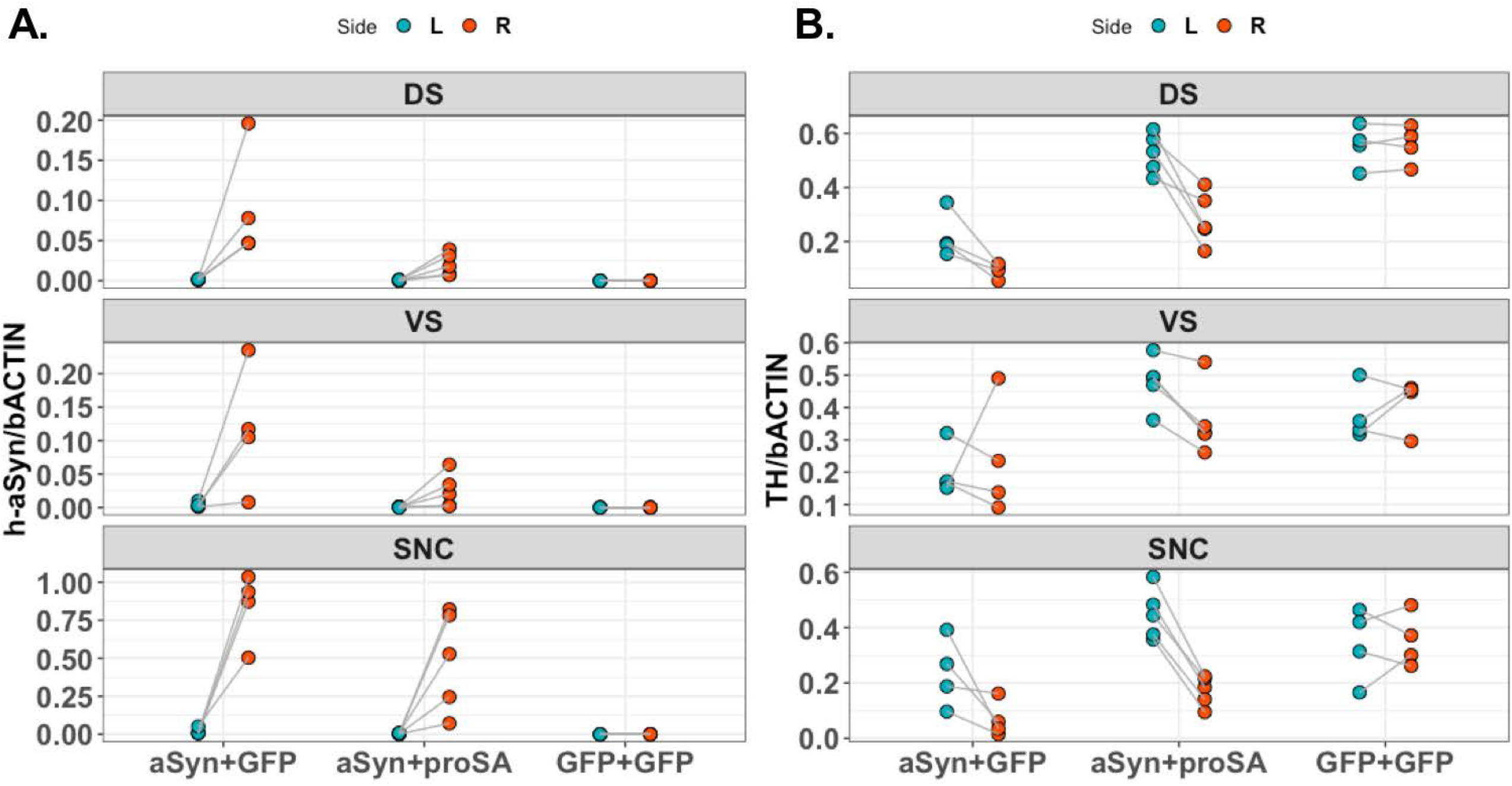
ProSAAS reduces human aSyn protein levels and attenuates the aSyn-mediated reduction in TH protein levels in nigrostriatal tissues. Bilateral quantitation of human aSyn and TH protein levels in the dorsal and ventral caudate and SNc at 3 weeks. Human aSyn expression was restricted to the injected side, as expected. TH protein levels were lower in aSyn + GFP-treated animals relative to GFP + GFP animals, an effect that, interestingly, was not restricted to the injected side. This effect of aSyn is attenuated bilaterally by unilateral proSAAS co-administration such that considerable asymmetry in TH levels persists.

